# Single-cell analysis identifies distinct CD4+ T cells associated with the pathobiology of pediatric obesity-related asthma

**DOI:** 10.1101/2024.08.13.607447

**Authors:** David A. Thompson, Yvonne B. Wabara, Sarai Duran, Anna Reichenbach, Laura Chen, Kayla Collado, Changsuek Yon, John M. Greally, Deepa Rastogi

## Abstract

Pediatric obesity-related asthma is characterized by non-atopic T helper 1 (Th1) inflammation and steroid resistance. CDC42 upregulation in CD4+T cells underliesTh1 inflammation but the CD4+T cell subtype(s) with CDC42 upregulation and their contribution to steroid resistance are not known. Compared to healthy-weight asthma, obesity-alone and healthy-weight controls, single-cell transcriptomics of obese asthma CD4+T cells revealed *CDC42* upregulation in 3 clusters comprised of naïve and central memory T cells, which differed from the cluster enriched for Th1 responses that was comprised of effector T cells. *NR3C1,* coding for glucocorticoid receptor, was downregulated, while genes coding for NLRP3 inflammasome were upregulated, in clusters with CDC42 upregulation and Th1 responses. Conserved genes in these clusters correlated with pulmonary function deficits in obese asthma. These findings suggest that several distinct CD4+T cell subtypes are programmed in obese asthma for CDC42 upregulation, Th1 inflammation, and steroid resistance, and together contribute to obese asthma phenotype.

**Summary:** CD4+T cells from obese children with asthma are distinctly programmed for non-allergic immune responses, steroid resistance and inflammasome activation, that underlie the obese asthma phenotype.

## Introduction

Asthma is one of the most common chronic diseases in children.^1^ Its pathobiology is heterogeneous and is broadly grouped into atopic and non-atopic asthma.^2^ Traditionally, the majority of pediatric asthma has been considered to be of atopic phenotype. However, there is evidence to suggest that up to 50% of pediatric asthma is non-atopic.^3^ Non-atopic asthma, best defined in the context of obesity-related asthma,^3^ is more severe with worse pulmonary function deficits, that are influenced by adiposity and non-atopic inflammation, rather than atopic sensitization.^4, 5, 6^ Obese asthma disease burden is poorly responsive to asthma medications, including inhaled and systemic steroids, which suggests steroid resistance.^7, 8^ Unlike the well-investigated pathobiology of atopic asthma that has informed therapeutics,^9^ the pathobiology of pediatric non-atopic asthma is poorly elucidated which has impeded development of targeted therapies.

Initial exploration of the pathobiology of obesity-related asthma has identified non-atopic systemic immune responses, with high CD4+T helper (Th) Th1/Th2 cell ratio and elevated non-atopic cytokines (tumor necrosis factor (TNF), interleukin (IL)-6, interferon (IFN)γ, and interferon-induced protein (IP)-10) in obese children with asthma that correlate with pulmonary function deficits.^10, 11^ Transcriptome-based investigation of mechanistic pathways underlying non-atopic immune responses in Th cells revealed upregulation of several genes in the Cell Division Cycle 42 (CDC42) pathway, which correlated with pulmonary function deficits in obesity-related asthma.^12, 13^ Small interfering RNA (siRNA)-mediated CDC42 silencing in CD4+T cells downregulated *IFNγ* and *TNF*, but not *IL-4,* expression verifying the role of CDC42 in Th1 polarization.^13^ Since CDC42 plays a seminal role in CD4+T cell chemotaxis and adhesion,^14, 15^ migration of obese asthma CD4+T cells with CDC42 upregulation was resistant to CDC42 inhibition, and these cells adhered more to airway smooth muscle (ASM) from obese donors, causing upregulation of pathways associated with ASM contractility and proliferation.^16^ These observations suggest that CD4+T cell with CDC42 upregulation is relevant to the pathobiology of non-atopic pediatric obesity-related asthma. Since steroids are more effective against atopic inflammation,^17^ we speculate that CDC42-mediated non-atopic Th1 inflammation is one mechanism that underlies steroid resistance in obese asthma. In addition, upregulation of NOD-like receptor family pyrin domain containing 3 (NLRP3) inflammasome, that activates IL-17 producing innate lymphoid cells and contributes to steroid resistance,^18^ has been reported in a murine model of obese asthma.^19^ While these investigations highlight the role of non-Th2 CD4+T cells in obese asthma pathobiology, the cellular characteristics of CD4+T cells that underlie CDC42 upregulation, steroid resistance, and inflammasome activation in pediatric obese asthma are not known. To address this gap in knowledge, we used single-cell transcriptomics to define the characteristics of CD4+T cell(s) with CDC42 upregulation, including their enrichment for non-Th2 or Th1 responses, steroid resistance and inflammasome activation,^18, 20, 21, 22, 23, 24^ and their association with pulmonary function deficits found in obesity-related asthma.^10, 11^

## Results

### Characteristics of the study cohort

Demographic characteristics did not differ between the four study groups in the full cohort **[Table 1]** or in the subset of 24 children (6 per group) that underwent CITE-Seq analysis **[Table S3]**. Children with obese asthma had lower FEV_1_/FVC ratio compared to all other groups and had higher IC compared to children with obesity-alone and healthy-weight controls in the full cohort; no differences were observed in the subset of 24 children.

**Table 1.** Demographic and clinical characteristics of study participants.

|  | **Obese asthma**<br>**(n=54)** | **Healthy-weight**<br>**asthma (n=27)** | **Obese no-**<br>**asthma (n=27)** | **Healthy-weight**<br>**controls (n=27)** | **Total** |
| --- | --- | --- | --- | --- | --- |
| **Demographics** |  |  |  |  |  |
| Age (years) | 11.56 ± 3.03 | 10.33 ± 3.17 | 11.11 ± 2.55 | 11.41 ± 3.12 | 0.37 |
| Male (n (%)) | 24 (44) | 15 (56) | 9 (33) | 15 (56) | 0.29 |
| African American (n (%)) | 38 (70) | 19 (70) | 19 (0.70) | 14 (52) | 0.42 |
| **Pulmonary Function Indices\*** |  |  |  |  |  |
| FVC | 101.54 ± 16.19 | 97.50 ± 17.74 | 103.16 ± 14.02 | 95.30 ± 13.18 | 0.21 |
| FEV<sub>1</sub> | 93.26 ± 19.40 | 93.73 ± 16.03 | 101.52 ± 11.67 | 94.70 ± 15.23 | 0.22 |
| FEV<sub>1</sub>/FVC ratio | 80.84 ± 8.84<sup>^</sup># | 85.85 ± 8.16<sup>^</sup> | 88.00 ± 5.00<sup>#</sup> | 88.07 ± 6.02<sup>#</sup> | <0.0001 |
| FEF<sub>25-75%</sub> | 79.88 ± 32.90 | 83.31 ± 27.92 | 96.76 ± 19.05 | 89.78 ± 19.88 | 0.07 |
| TLC | 94.92 ± 15.58 | 94.58 ± 20.31 | 97.50 ± 23.76 | 94.54 ± 12.51 | 0.93 |
| RV | 105.53 ± 42.71 | 123 ± 47.18 | 108.82 ± 86.04 | 106.25 ± 30.69 | 0.60 |
| RV/TLC ratio | 24.12 ± 7.26 | 27.96 ± 8.56 | 22.91 ± 10.40 | 24.38 ± 5.70 | 0.16 |
| ERV | 67.35 ± 64.24 | 77.29 ± 40.30 | 68.16 ± 27.63 | 81.93 ± 26.76 | 0.55 |
| FRC | 86.04 ± 22.33 | 96.67 ± 24.92 | 89.64 ± 44.14 | 96.88 ± 17.08 | 0.29 |
| IC | 102.33 ± 20.94<sup>^</sup> | 90.64 ± 24.68 | 103.44 ± 14.49 | 89.33 ± 16.41<sup>^</sup> | 0.007 |
| Atopic sensitization<br>(n (%)) | 3 (5.5) | 4 (14.8) | 0 (0) | 3 (11.1) | 0.16 |

\* Percent predicted values of pulmonary function indices other than FEV<sub>1</sub>/FVC ratio and RV/TLC ratio are reported.
Between-group comparisons were done using Analysis of Variance with post-hoc Bonferroni correction for continuous variables and $\chi^2$ test for categorical variables. The significance of between-group differences is noted as follows: <sup>^</sup> p<0.05 <sup>#</sup> p<0.01

### sc-RNA-seq based analysis of CD4+ cells from obese asthma, healthy-weight asthma, obesity-alone or healthy-control groups

CITE-seq analysis of CD4+T cells combined for all study groups identified 22 cell clusters (clusters 0 to 21) **[Fig. 1a].** Azimuth-based cell classification^25^ across samples verified 95-99.9% of sequenced cells were CD4+T cells, and included naïve T cells, three subsets of αβ central memory T (TCM) cells, three subsets of αβ effector memory (TEM), four subsets of γδ (gdT) T cells, naïve and memory T regulatory (Tregs) cells, and two subsets of double negative (dnT) T cells. The remaining 0.01-5% of cells included cytotoxic lymphocyteheatmaps summarize the association of percents (CTL), five subsets of natural killer (NK) cells, innate lymphoid cells (ILCs), plasmacytoid, transitional, and two subsets of conventional dendritic cells (pDCs, ASDCs, and cDCs), mucosal-associated invariant T (MAIT) cells, plasma cells, hematopoetic stem and progenitor cells (HSPC), and CD14 and CD16 monocytes for all but two participants, one with obese asthma and one with obesity-alone, where these cells subtypes comprised 6 to 10% cells **[Fig. 1b, Table S4]**. There was lack of complete overlap between clusters and Azimuth-based cell classification such that most clusters were comprised of more than two Azimuth cell subtypes **[Fig. 1c, Table S4]**. Using transcription factors to classify clusters into Th cell subsets, we identified enrichment of the *TBX21* transcription factor for Th1 cells in cluster 12 and of the *FOXP3* transcription factor for Tregs in cluster 10. Other transcription factors were enriched in multiple clusters, including the *GATA3* transcription factor for Th2 cells enriched in clusters 5, 6, 10, 12, 15, 17, and 19, and relative enrichment, despite limited expression, of the *RORC* transcription factor for Th17 cells in clusters 5, 9, 10 and 15 **[Fig. 1d]**. Overlap with a published reference set for CD4+ T cell subsets^26^ verified these classifications **[Fig. 1e]**, and in keeping with failure of Azimuth and transcription factor approach to classify the majority of clusters into a single known CD4+T cell subtype, highlighted CD4+T cell heterogeneity.

**Figure 1.**
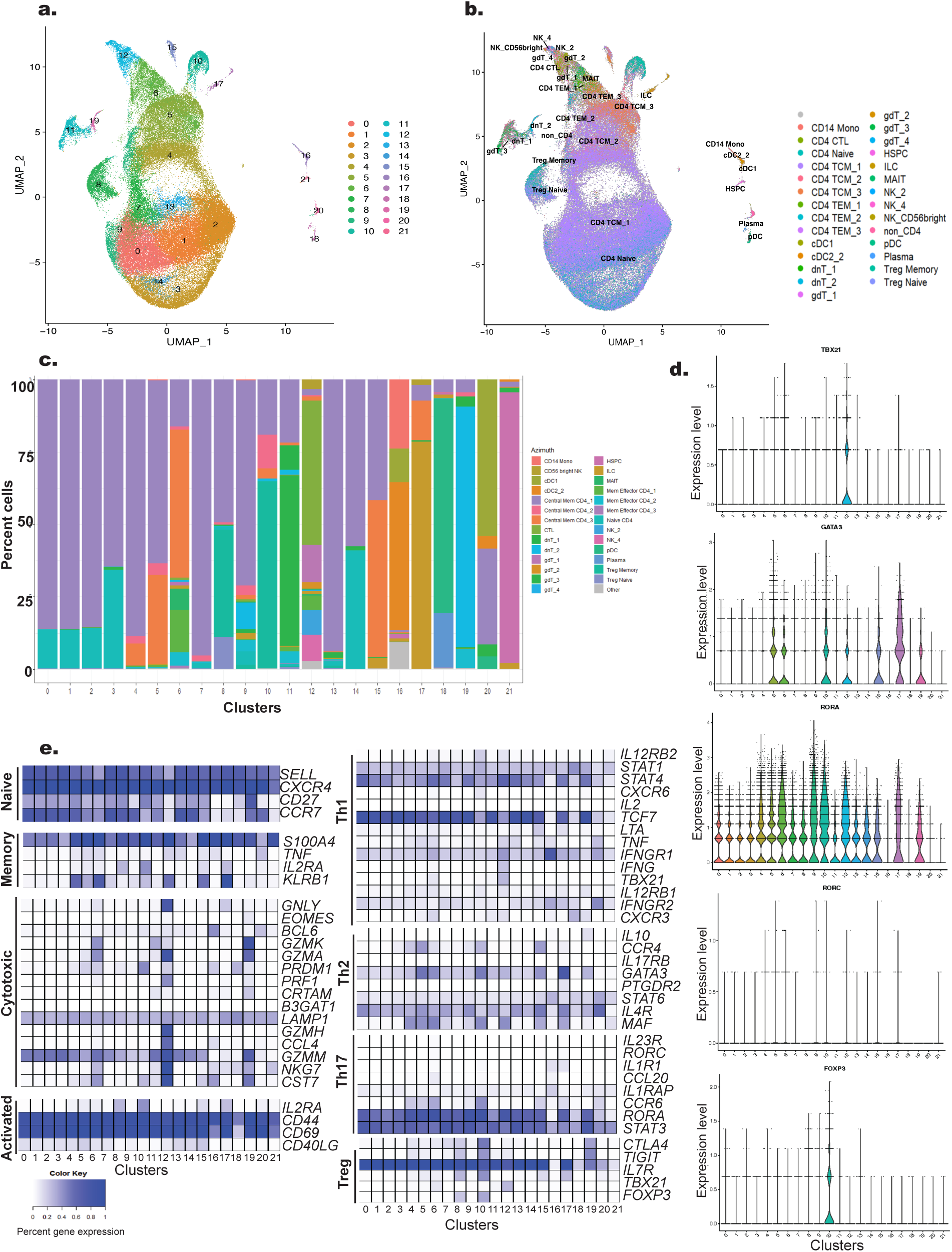
CITE-Seq analysis. **a)** UMAP plot of CITE-Seq analysis revealed 22 clusters (0 to 21) within CD4+T cells from all four study groups. **b)** Azimuth-based classification of CD4+T cells revealed CD4+T cell subtypes do not completely overlap with the 22 clusters. **c)** Stacked bar plot summarizes the proportions of Azimuth-based CD4+T cell subtypes in each cluster. **d)** Cluster-specific expression of Th cell subset-specific transcription factors reveal generalized lack of any cluster for a specific Th cell subset other than cluster 12 for Th1 and cluster 10 for T regulatory cells. **e)** Azimuth-based T cell classification overlapped to large extent with those published by Ding et al.^26^

### Identification of clusters with CDC42 upregulation in obese asthma

Investigating for clusters enriched for CDC42 in obese asthma, we found that *CDC42* expression was higher in obese asthma in clusters 0, 3, and 8 compared to healthy-weight asthma, obesity-alone, and healthy controls **[Fig. 2a, b]**. Azimuth-based classification revealed that clusters 0 and 3 were primarily comprised of αβ TCM (86% and 64.5% respectively) and naïve T cells (13% and 33% respectively), while cluster 8 was comprised of 49% αβ TCM cells, and 37% memory and 10% naïve Tregs **[Fig. 2c]**. There were significantly fewer cells in cluster 3 in obese asthma **[Fig. 2d]** but this difference was not driven by any specific Azimuth subtype. Higher *CDC42* expression in obese asthma was present in naïve and αβ TCM cells and memory Tregs but not in naïve T regs **[Fig. 2e]**. Between-group comparison of additional members of the CDC42 pathway including guanine exchange factors upstream of CDC42, Dedicator Of Cytokinesis 5 *(DOCK5)* and vav guanine nucleotide exchange factor 2 *(VAV2),* and kinases downstream of CDC42, including p21 activated kinase 3 *(PAK3)*, mitogen activated protein kinase kinase kinase 11 *(MAP3K11 or MLK3),* and nuclear factor kappa B *(NFKB1)*, revealed enrichment of *DOCK5* in cluster 8, while *NFKB1* was enriched in clusters 0 and 3 in obese asthma, particularly compared to healthy-weight asthma **[Fig. 2f]**. While *NFKB1* was enriched in obese asthma compared to other study groups in naïve and αβ TCM cells **[Fig. 2g]**, *DOCK5* was not enriched in any specific Azimuth cell subtype. In keeping with classification of cells as naïve and central memory T cells, no transcription factor associated with effector Th cells (*TBX21*, *GATA3*, or *RORC*) was enriched in clusters 0, 3, and 8 **[Fig. 2h]**.

**Figure 2.**
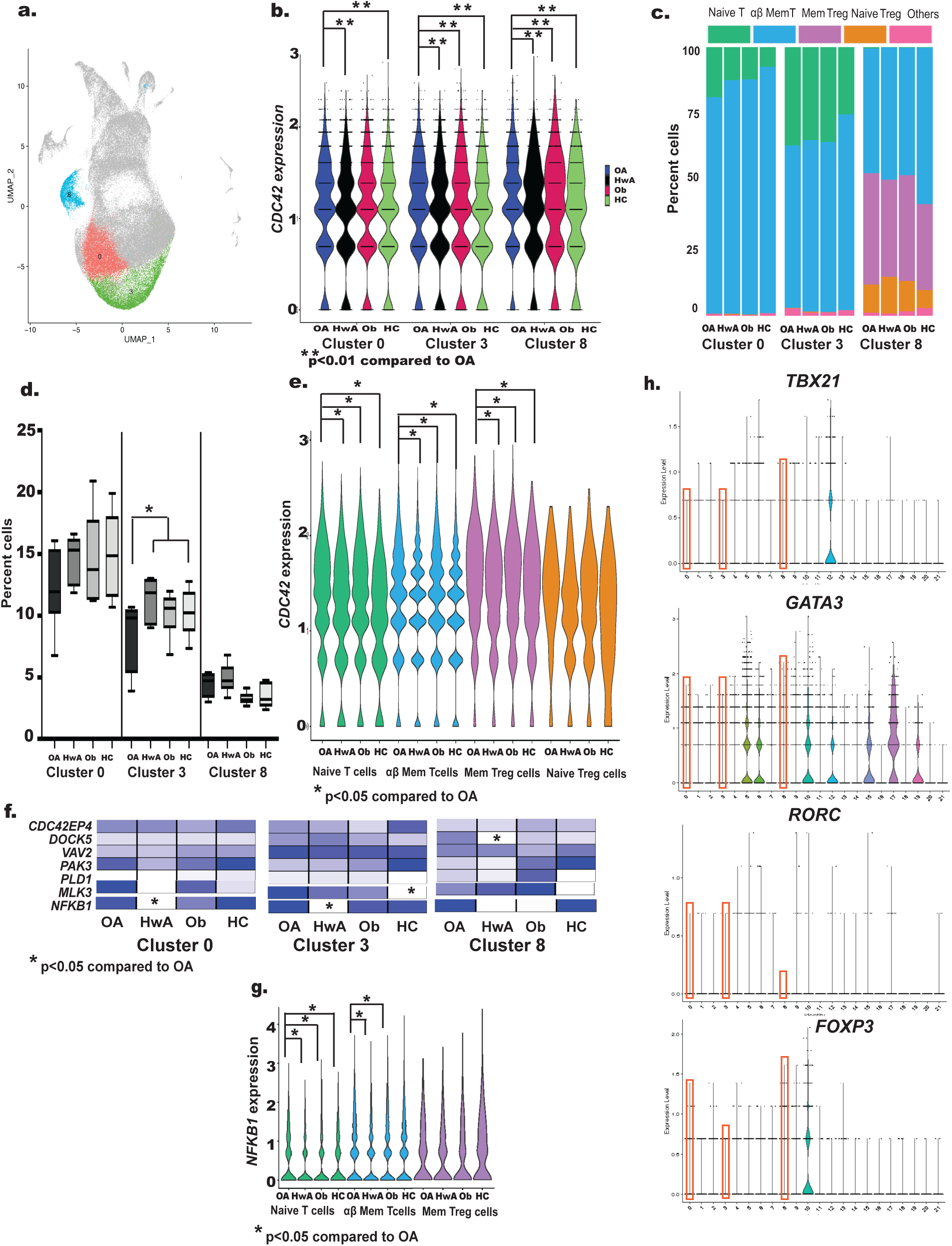
Identification of cell clusters with *CDC42* enrichment in obese asthma. **a)** The three clusters (0, 3, and 8) enriched for CDC42 in obese asthma are marked on the cell cluster UMAP. **b)** Comparison of *CDC42* expression between the four study groups (obese asthma (OA), healthy-weight asthma (HwA), obese no-asthma (Ob) and healthy-weight controls (HC) in clusters 0, 3, and 8 revealed its significant upregulation in OA in all three clusters. **c)** Stacked bar plot summarizes the proportion of Azimuth-based CD4+T cell subtypes in clusters 0, 3, and 8. **d)** Between study-group comparisons of cell proportions in each cluster reveal significantly fewer cells in OA in Cluster 3 as compared to the other study groups. **e)** Between-study group comparison of *CDC42* expression in Azimuth-based CD4+T cell subtypes in clusters 0, 3, and 8 suggest that CDC42 is enriched in naïve, αβ central memory T cells, and memory T regulatory cells but not in naïve T regulatory cells. **f)** Between-study-group comparison of additional genes in the CDC42 pathway in clusters 0, 3, and 8.^13^ Higher intensity of blue denotes higher expression of the specific gene in each study group. Absent expression is depicted in white. *NFKB1* expression in Clusters 0 and 3 and *DOCK5* expression in Cluster 8 were significantly higher in OA compared to HwA. **g)** Among CD4+T cell subtypes enriched for *CDC42* expression, *NFKB1* expression was higher in OA compared to the other groups in naïve T cells and in OA compared to HwA and Ob in αβ central memory T cells. h) None of the known Th cell transcription factors was enriched in clusters 0, 3, and 8.

Since CDC42 was enriched in three clusters in obese asthma, we investigated for overlap in conserved genes between these clusters. We found 20 genes, including 13 conserved genes for cluster 0, 3 conserved genes for cluster 3, and 4 conserved genes for cluster 8 that were differentially expressed in obese asthma compared to other groups in all three clusters **[Fig 3a]**. Of these, 10 were downregulated while 10 were upregulated in obese asthma compared to other groups in all 3 clusters **[Fig. 3a]**. Network analysis of the 20 genes **[Fig. 3b]** revealed enrichment for Th1 and Th2 differentiation, Primary Immunodeficiency, Hematopoetic cell lineage, and T cell receptor signaling pathway in Kyoto Encyclopedia of Genes and Genomes (KEGG) pathways, and RNA polymerase II transcription regulator complex in Gene Ontology (GO) Cellular Compartment analysis, suggesting redundancy of these CD4+T cellular processes in clusters 0, 3, and 8.

**Figure 3.**
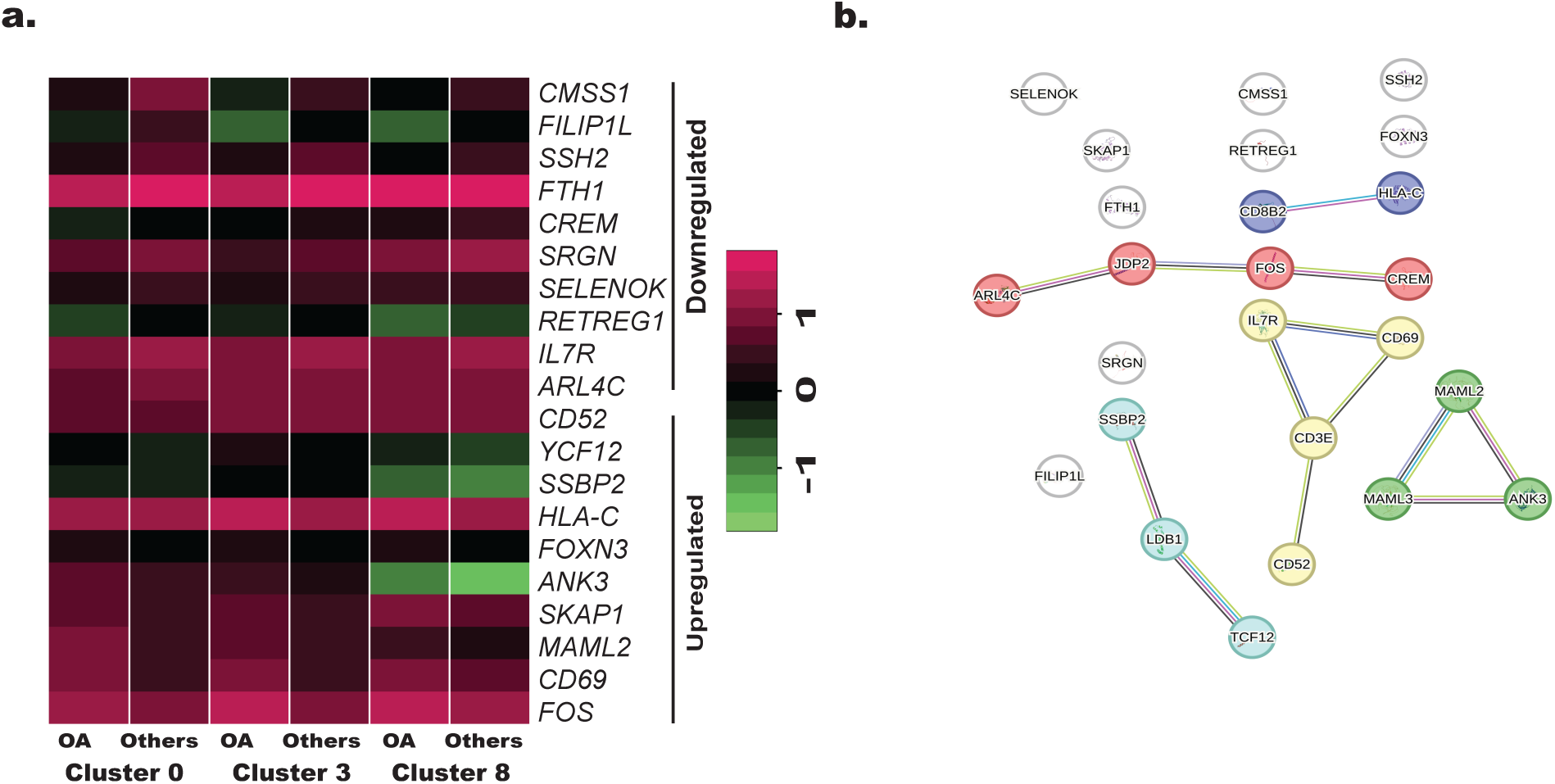
**a)** Heatmap of the 20 conserved genes in clusters 0, 3, and 8 that were up- or downregulated in obese asthma (OA) as compared to other three groups (Others)**. b)** KEGG pathway analysis of these 20 genes revealed enrichment for Th1 and Th2 differentiation, Primary Immunodeficiency, Hematopoetic cell lineage, and T cell receptor signaling pathway.

### Identification of clusters associated with Th1 polarization in obese asthma

Since CDC42 enrichment in clusters 0, 3, and 8 did not overlap with Th1 polarization, we investigated other clusters and found enrichment of the *TBX21* transcription factor for Th1 cells, and downstream Th1 cytokines, *IFNG* and *TNF,* uniquely in cluster 12 **[Fig. 4a-c]**. Cell proportions in cluster 12 were higher in obese asthma compared to other groups **[Fig. 4d],** and inversely correlated with proportions in cluster 3 only in obese asthma **[Fig. 4e]**. Based on Azimuth classification, cluster 12 was enriched for several types of effector T cells, including cytotoxic T cells, two subsets of γδ T cells, a subset of αβ effector memory T cells and NK cells in addition to 2 subsets of central memory T cells **[Fig. 4f]**. The between-group differences in cell proportions in cluster 12 were not explained by any of these cell subtypes. Given the diverse cell subtypes, we investigated differential enrichment of additional transcription factors in cluster 12. Although *TBX21* was enriched in cluster 12 **[Fig. 4a]**, its expression in obese asthma did not differ from the other groups **[Fig. 4g],** a pattern that was also observed for *IFNG* and *TNF* **[Fig. 4h, i]**. Conversely, while the *RORA* transcription factor for Th17 cells was not enriched in cluster 12, its expression was significantly higher in obese asthma compared to other groups **[Fig. 4j, k];** no differences were observed for *GATA3, RORC*, or *FOXP3* **[Fig. 1d]**. To elucidate conserved genes in cluster 12 that may underlie Th1 polarization and be enriched in obese asthma, we identified 27 conserved genes **[Table. S5]** that were differentially expressed in obese asthma compared to other groups. Network analysis of these 27 genes revealed enrichment for Th1, Th2, and Th17 differentiation, cytokine-cytokine receptor interaction, graft vs. host disease, apoptosis, and phagosome pathway in KEGG pathways, and immune response in GO Cellular Processes **[Fig. 4l]**.

**Figure 4.**
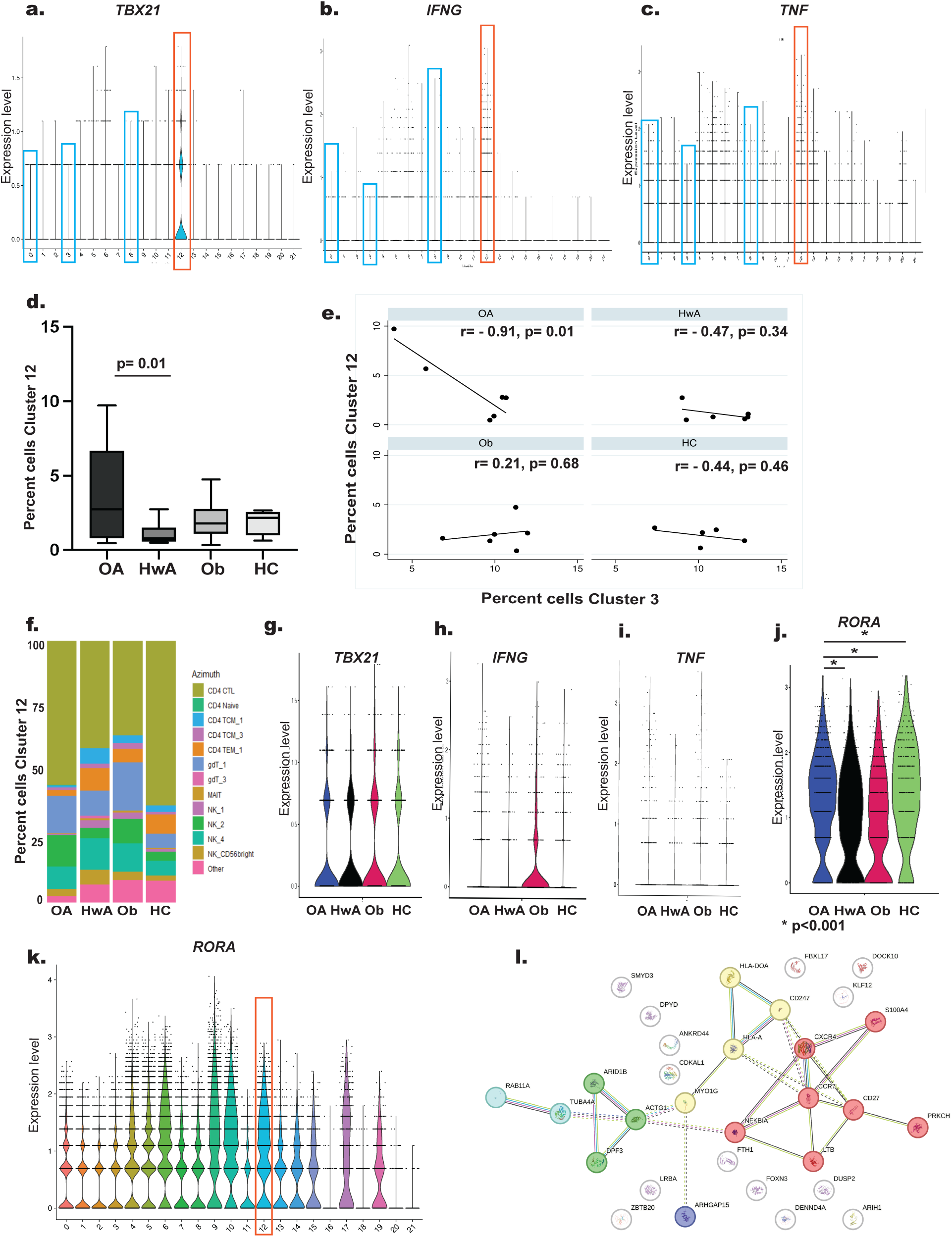
**a)** Expression of the *TBX21* transcription factor for Th1 cells was enriched in Cluster 12, as compared to Clusters 0, 3, and 8. There was corresponding enrichment of Th1 cytokines **b)** *IFNG* and **c)** *TNF* in Cluster 12. **d)** Comparisons of cell proportions in Cluster 12 between study-groups (obese asthma (OA), healthy-weight asthma (HwA), obese no-asthma (Ob) and healthy-weight controls (HC) revealed significantly higher proportion of cells in OA in Cluster 12. **e)** Study-group specific correlations between cellular proportions in clusters 3 and 12 revealed significant correlation only in OA. **f)** Stacked bar plot summarizing study-group specific distribution of Azimuth-based CD4+T cell subtypes revealed several different effector cells enriched in cluster 12. Comparison of **g)** *TBX21* expression, **h)** *IFNG* expression, and **i)** *TNF* expression showed absence of differences between study groups. **j)** Between-study group comparison of *RORA* expression revealed its enrichment in OA compared to other three groups. **k)** Violin plot of *RORA* expression in all clusters revealed that it was not enriched in Cluster 12 as compared to other clusters. **l)** KEGG pathway analysis of 27 conserved genes in cluster 12 that were differentially regulated in OA as compared the other three groups revealed enrichment for Th1, Th2, and Th17 differentiation, cytokine-cytokine receptor interaction, graft vs. host disease, apoptosis, and phagosome pathway.

### Identification of clusters with enrichment for genes associated with steroid resistance in asthma

Given presence of steroid resistance and inflammasome activation in obese asthma,^7, 19^ we next investigated the overlap of CDC42 upregulation and Th1 polarization with steroid resistance and inflammasome activation. We found that clusters 0, 3, and 8 with CDC42 upregulation and cluster 12 with Th1 polarization were also enriched for steroid resistance and inflammasome activation, as were additional clusters 1, 2, 4, 5, 6, 11, 16 and 18. From the perspective of steroid resistance, nuclear receptor subfamily 3 group C member 1 (*NR3C1),* the gene encoding for glucocorticoid receptor, was downregulated in obese asthma compared to other groups in clusters 0, 1, 2, 3, 4, 5, 6, 8, 11 and 12 **[Fig. 5a]**, and its downstream target, *NFKB1,* was upregulated in all these clusters except cluster 1 **[Fig. 5b]**. From the perspective of inflammasome activation, phosphatase and tensin homolog (*PTEN)* was upregulated in clusters 0, 1, 2, 3, 8, 11, and 12 **[Fig. 5c]**, caspase 1 *(CASP1)* **[Fig. 5d]** was upregulated in clusters 1, 4, 5, and 6, and its downstream cytokine, *IL1B,* was upregulated in cluster 16 **[Fig. 5e]**. Azimuth classification revealed clusters 1, 2, 4, 5, and 6 were enriched for αβ central memory T cells, cluster 11 was enriched for γδ T cells, cluster 16 was enriched for CD14 monocytes and classical DCs, and cluster 18 was enriched for plasmacytoid DCs and plasma cells **[Fig. 5f]**. Cell proportions were higher in obese asthma group in cluster 16 as compared to healthy-weight asthma but did not differ from obesity-alone or healthy-control in cluster 16 **[Fig. 5g]**; there was no difference in cell proportions in obese asthma and other groups in any of the other clusters.

**Figure 5.**
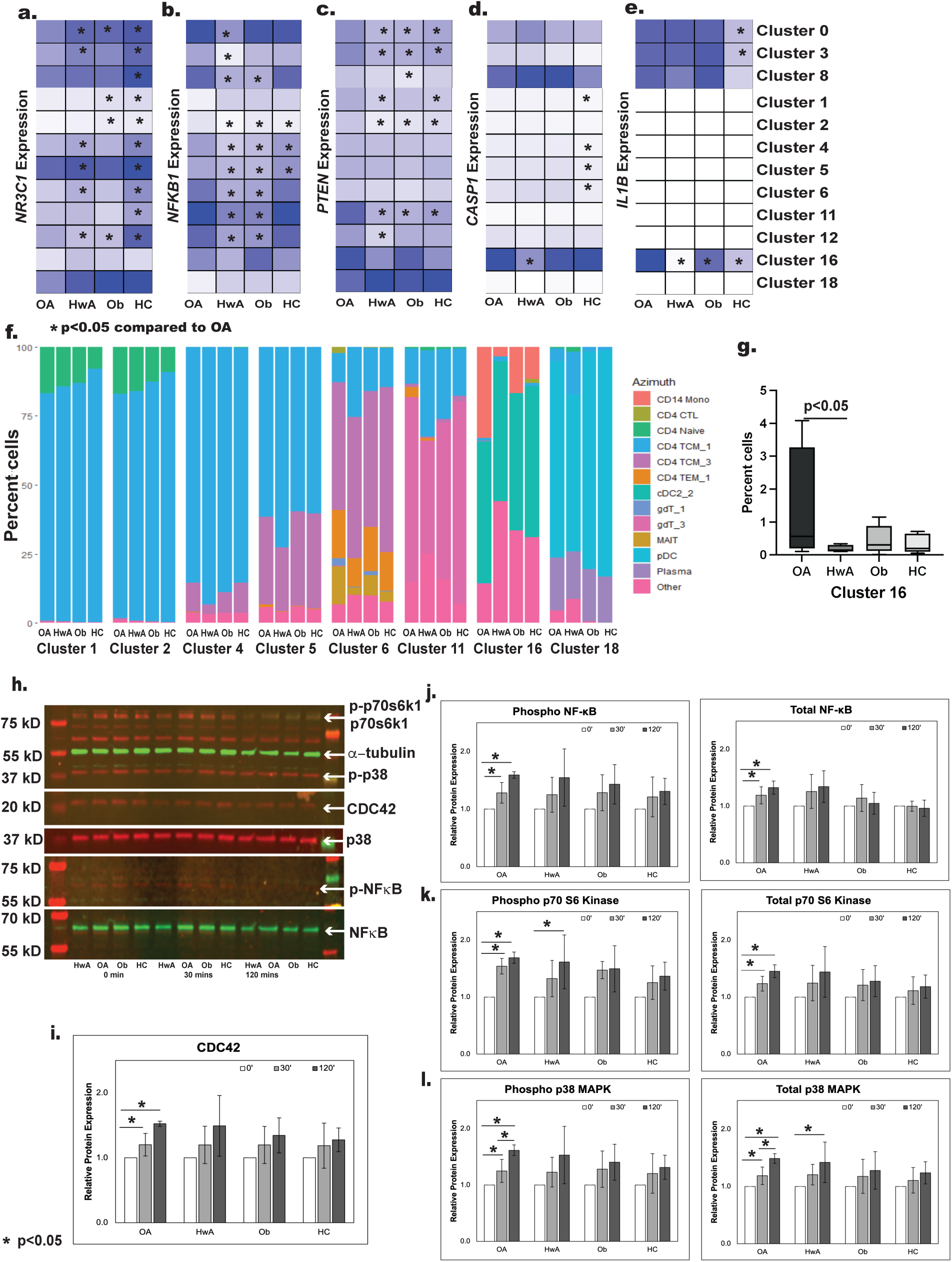
Investigation of steroid resistance and inflammasome activation between obese asthma (OA) and other three groups (healthy-weight asthma (HwA), obese no-asthma (Ob) and healthy-weight controls (HC)) revealed **a)** downregulation of *NR3C1*, the gene coding for glucocorticoid receptor in clusters 0, 1, 2, 3, 4, 5, 6, 8, 11 and 12**, b)** upregulation of *NFKB1* in clusters 0, 2, 3, 4, 5, 6, 8, 11, and 12 and, **c)** upregulation of *PTEN,* associated with inflammasome activation, in clusters 0, 1, 2, 3, 8, 11, and 12. Downstream of *PTEN*, **d)** *CASP1* was upregulated in OA in clusters 1, 4, 5, and 6, while **e)** *IL1B* was upregulated specifically in cluster 16. Higher intensity of blue denotes higher expression of the specific gene in each study group. Absent expression is depicted in white. Statistically significant differences between OA and HwA, Ob and HC are denoted by the asterisk. **f)** Stacked bar plot summarizes proportion of Azimuth-based CD4+ cell subtypes in clusters 1, 2, 4, 5, 6, 11, 16 and 18. **g)** Between-study group comparison of cell proportions in these clusters were only different between OA and other study groups in cluster 16 where cell proportions were the highest in OA. **h)** Protein bands from a representative blot of the 5 replicates is shown here. Each blot included protein lysates collected at 30 minutes and 2 hours post TCR stimulation from one sample from each study group. Protein levels were quantified using the LICOR system, where secondary antibody to rabbit primary antibodies (CDC42, p-NFκB, p38 MAPK, p-p38 MAPK, and p70s6k1) fluoresced at 780nm (red) and secondary antibody to mouse primary antibodies (total NFκB, p-p70s6k1, and α-tubulin) fluoresced at 680nm (green). Protein levels were normalized to α-tubulin. **i)** CDC42, **j)** phospho- and total NFκB, **k)** phospho- and total p70s6k1, and **l)** phospho- and total p38 MAPK were increased at 30 minutes and 2 hours in OA samples. In addition, phospho-p70s6k1 and total p38 MAPK were also increased in HwA at 2 hours but not at 30 minutes.

### Protein-based verification of differential gene expression in obese asthma

As verification of *CDC42* and *NFKB1* enrichment in obese asthma, protein levels of CDC42, and total and phospho-NFκB were increased only in obese asthma after 30-minute and 2-hour T cell receptor (TCR) stimulation **[Fig. 5h-j]**. Elucidation of differential expression of signaling pathways downstream of CDC42 that are associated with NFκB activation in T cells revealed upregulation of phospho- and total p70s6k1 and phospho- and total p38 MAPK in obese asthma at 30 minutes and 2 hours, while phopho-p70s6k1 and total p38MAPK was upregulated in healthy-weight asthma at 2 hours but not at 30 minutes **[Fig. 5k,l]**. Neither phospho- nor total p70s6k1 or p38MAPK were increased in obesity-alone or healthy-weight controls after 30-minute and 2-hour TCR stimulation.

Clinical relevance of genes associated with CDC42 upregulation, Th1 polarization, and steroid resistance.

To investigate the clinical relevance of genes differentially enriched in clusters 0, 3, 8 and 12, and those associated with steroid resistance, in obese asthma as compared to other three groups, we quantified the association of their CD4+T cell expression measured with bulk RNA Seq in a larger cohort of 135 participants with FEV_1_/FVC ratio and percent predicted ERV, the two most frequently indices affected in obese asthma. We limited this investigation to obese asthma as compared to healthy-weight asthma due to clinical relevance of pulmonary function in the context of asthma. Overall, as compared to 5% genes correlating with FEV_1_/FVC ratio and percent predicted ERV by chance, 18% of conserved genes in cluster 8 were significantly correlated with FEV_1_/FVC ratio in obese asthma and 20% of conserved genes in clusters 3 and 8 correlated with ERV in healthy-weight asthma. Among the 20 genes conserved in clusters 0, 3, and 8, reticulophagy regulator 1 (*RETREG1)*, and interleukin 7 receptor (*IL7R)*, were upregulated, and among 27 genes conserved in cluster 12, dihydropyrimidine dehydrogenase *(DPYD)* and LPS responsive beige-like anchor protein *(LRBA)* were upregulated in obese asthma as compared to other groups, and inversely correlated with FEV_1_/FVC ratio **[Fig. 6a, b]**. Among the inflammasome genes, *CASP1*, upregulated in obese asthma, was inversely correlated with FEV_1_/FVC ratio, while *NR3C1*, downregulated in obese asthma, directly correlated with ERV **[Fig. 6c]**.

**Figure 6.**
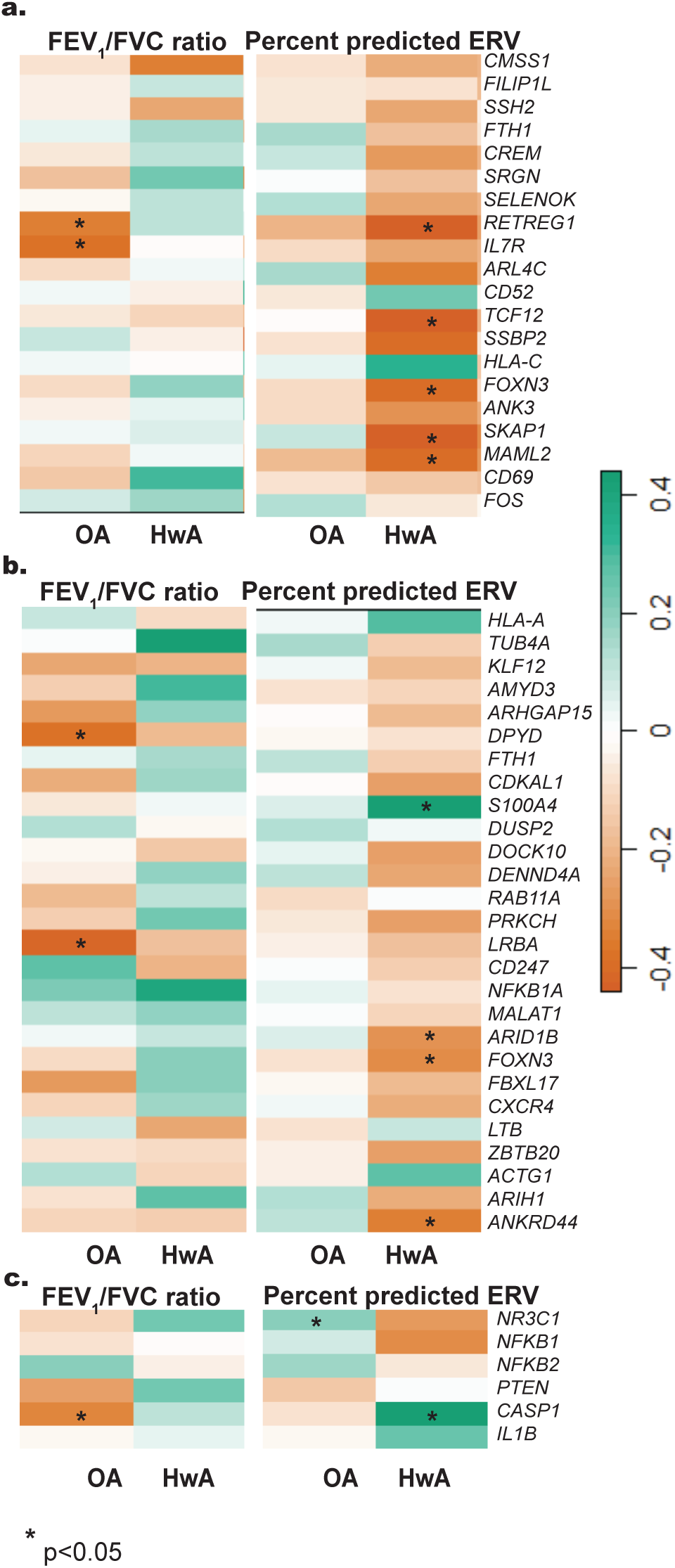
These heatmaps summarize the association of percent FEV_1_/FVC ratio and percent predicted ERV in obese asthma (OA) and healthy-weight asthma (HwA) with **a)** 20 conserved genes in clusters 0,3, and 8 that were up- or downregulated in OA as compared to other three groups, **b)** 27 conserved genes in cluster 12 differentially expressed in OA as compared to the other three groups, and **c)** 6 genes that are markers of steroid resistance or inflammasome activation and were differentially expressed in OA as compared to other three groups. Expression of genes that significantly correlated with lung function indices in OA or HwA are marked with an asterisk. The color key summarizes the range of r values of the correlation and their directionality.

## Discussion

Using single-cell RNA sequencing, we report for the first time on CD4+T cell programming that underlies non-atopic T cell responses, including Th1 polarization with CDC42 upregulation, in pediatric obesity-related asthma.^12, 13^ In keeping with the heterogeneity of CD4+T cells, we identified 22 distinct clusters, of which three clusters comprised of naïve and αβ central memory CD4+T cells and memory T regulatory cells were enriched for CDC42 pathway in obese asthma compared to healthy-weight asthma, obesity-alone and healthy-weight controls. A fourth distinct cluster, comprised of several effector cell subtypes including cytotoxic T cells, αβ central memory, αβ effector memory, and γδ T cells, and NK cells was enriched for Th1 polarization and had the highest proportion of cells in obese asthma samples. *NR3C1*, the gene encoding for the glucocorticoid receptor, was downregulated, and *NFKB1* as well as markers of inflammasome activation were upregulated, in the three clusters with CDC42 upregulation as well as the one with Th1 polarization, directly linking CDC42 enrichment and Th1 polarization in obese asthma with inflammasome activation and steroid resistance.^20^ Elevated protein levels of CDC42 and NFκB in obese asthma samples distinct from those that underwent sc-RNA Seq analysis verified the transcriptomic findings. Moreover, *NFKB* expression was enriched in 5 additional clusters, which were also comprised of naïve, αβ central memory, effector memory, and γδ CD4+T cells suggesting that there is programming of additional CD4+T cells for non-atopic pro-inflammatory responses. Lastly, several conserved genes in the three clusters with CDC42 upregulation and in the cluster with Th1 polarization as well as those associated with steroid resistance correlated with pulmonary function deficits verifying their clinical relevance in the obese asthma phenotype.

Given that CDC42 upregulation in obese asthma occurs in cells distinct from those with Th1 polarization, but both are enriched for steroid resistance and the inflammasome and correlate with the obese asthma phenotype, suggest that several subtypes of CD4+T cells, rather than a single subtype, are differentially programmed in obese asthma and link with its phenotype.

Enrichment of the three clusters with CDC42 upregulation for naïve and αβ central memory CD4+T cells suggests that obese asthma is associated with long-term CD4+T cell programming capable of inducing rapid effector T cell responses.^27^ Despite enrichment for *CDC42*, cell proportions in the one cluster (cluster 3) that had the highest proportion of naïve CD4+T cells, were lower in obese asthma, suggesting that naïve T cells are decreased in number but facilitate the pro-inflammatory state of obese asthma. This speculation was supported by the upregulation of *NFKB1* in both naïve and αβ central memory CD4+T cells in obese asthma, and the inverse correlation between cell proportions in cluster 3 with proportions of cluster 12, the cluster with Th1 polarization, wherein fewer cells in cluster with CDC42 upregulation directly correlated with higher proportion of cells with Th1 polarization, uniquely in obese asthma. These differences in cluster-specific gene expression as well as in cell proportions highlights the importance of qualitative as well as quantitative changes in cells that can be elucidated by sc-RNA-Seq.^28^ In keeping with the overlap of gene expression and Azimuth-based cell subtypes between the three clusters with CDC42 upregulation, 20 conserved genes in these clusters were consistently up- or downregulated in all three clusters in obese asthma relative to other three groups. Upregulated genes included several T cell activation genes such as *CD69* and *CD52*, associated with T cell activation and proliferation,^29, 30^ *FOS*, a subunit of AP-1 complex that is pro-inflammatory,^31^ *SKAP1*, Src Kinase-Associated Phosphoprotein 1, a T cell adaptor protein, that positively regulates T-cell receptor signaling and migration,^32^ and *TCF12*, transcription factor 12, that causes lineage-specific transition from double negative to double positive T cells.^33^ Conversely, genes that were downregulated have inhibitory effects on T cell activation and function. For example, *FTH1*, that encodes for heavy chain ferritin, decreases T cell inflammation by activating T regulatory cells,^34^ *SSH2,* slingshot protein phosphatase 2, a regulator of actin polymerization,^35^ controls T cell migration, *IL7R*, plays a critical role in lymphocyte development, interacts with CDC42 for T cell survival, and is downregulated with T cell receptor activation,^36^ and *SELENOK*, selenoprotein K, controls T-cell proliferation and migration.^37^ The directionality of gene expression in both up- and downregulated genes cumulatively was in favor of T cell activation, which was verified by the enrichment of Th1 and Th2 differentiation and T cell receptor signaling in network analysis of these genes suggesting that these clusters, although enriched in naïve and central memory T cells, are intricately linked with Th cell differentiation and activation in obese asthma.

The cluster enriched for Th1 polarization was distinct from the clusters enriched for CDC42, had higher proportion of cells in obese asthma as compared to the other three groups, and was comprised of several types of effector T cells, including cytotoxic T cells, αβ central memory, αβ effector memory, and γδ T cells, and NK cells. No specific CD4+T cell subtype underlay Th1 polarization or higher proportion of cells in this cluster in obese asthma. In keeping with the importance of qualitative and quantitative differences, while expression of *TBX21*, that encodes for T-bet transcription factor for Th1 cells, did not differ between obese asthma and other groups, *RORA* was most upregulated in obese asthma compared to the other three groups. *RORA* encodes for RORα that functions as a transcription factor and facilitates differentiation of ILCs to ILC3 and of naïve Th cells to Th17 cells, typically in the absence of or concurrently with RORγt, the classical transcription factor for Th17 cells.^38, 39^ Higher Th17 cells and IL-17 concentrations in serum and airway samples have been linked with severe asthma that is steroid resistant, likely due to neutrophil-dominant inflammation.^40, 41^ Our observations therefore suggest that augmented Th17 responses in presence of Th1 polarization in obese asthma likely distinguish it from the other three groups and underlie disease severity and steroid resistance.^7, 8^ Our speculations are verified by the contribution of IL-17 to steroid resistance in obese asthma by upregulation of glucocorticoid receptor-beta (GR-β), a dominant-negative inhibitor of GR-α, the classical active receptor for steroids,^42^ in adipocytes and serum from obese adults with asthma.^43^ *RORA* upregulation is also of high relevance in our study because it is intricately linked to HIF1α,^44^ a key player in Th cell immunometabolism. Upon activation, Th cells transition from oxidative phosphorylation to glycolysis.^45^ Hyperglycemia, one of the integral components of insulin resistance, facilitates Th1 cell activation by activating PI3k/Akt and downstream mTORC1 and HIF1α signaling,^46^ which increases glycolysis.^47^ In light of the known association of insulin resistance with obese asthma disease burden and Th1 polarization,^4, 11^ *RORA* upregulation identifies HIF1α as a potential molecular link between insulin resistance, Th1 polarization, and pulmonary function deficits found in obese asthma.

The three clusters with CDC42 upregulation and the one with Th1 polarization had evidence of steroid resistance with *NR3C1* downregulation and inflammasome activation with *PTEN* upregulation.^48^ These observations extend the known link of CDC42 upregulation and non-T2 immune responses with steroid resistance and inflammasome activation^49, 50^ to the pathobiology of obesity-related asthma. *NR3C1* downregulation was observed in seven additional clusters other than those with *CDC42* upregulation and Th1 polarization and *NFKB1* and/or *PTEN* was upregulated in a subset of these clusters, suggesting that steroid resistance in obese asthma is not driven by any one or a few CD4+T cells subtypes but is due to generalized differential programming of CD4+T cell transcriptome in obese asthma as compared to healthy-weight asthma, obese alone or healthy controls. We verified these observations by demonstrating higher levels of total NFκB protein levels at 30 minutes and 2 hours after T cell stimulation in obese asthma compared to other three groups. Furthermore, in keeping with PTEN within the NRLP3 inflammasome pathway causing upregulation of IL-1β,^48^ we found higher IL-1β in obese asthma that was driven exclusively by a cluster comprised of CD14+ and CD16+ monocytes. In light of prior reports of higher patrolling (CD14loCD16+) or activated monocytes in adolescents with obese asthma,^11^ our findings link CD4+T cell responses with monocyte activation in obese asthma via inflammasome activation in CD4+T cells. These cellular mechanisms also likely underlie activation of the inflammasome reported in obese adults with asthma in response to diets enriched in saturated fats,^51^ and its attenuation by involucrasin B, an isoflavinoid found in plants, in a murine model of obese asthma,^52^ and are targets of future investigations.

The functional relevance of the CD4+ T cell transcriptome in obese asthma was further supported by the enrichment between gene expression and FEV_1_/FVC ratio in obese asthma. *RETREG1* and *IL7R* among genes in the clusters with CDC42 upregulation, and *DPYD* and *LRBA* in the cluster with Th1 polarization, and *CASP1*, among genes associated with the inflammasome, link pro-inflammatory T cell activation and infiltration with the obese asthma phenotype. Intriguingly, *NR3C1*, the gene coding for the glucocorticoid receptor, was the only gene that directly correlated with ERV, suggesting its downregulation correlated with lower ERV in obese asthma, further linking steroid resistance with lower ERV, a consistent and specific feature of obese asthma.^53^

Bringing all our observations together, using single-cell sequencing, we report on differential programming of obese asthma CD4+T cells as compared to healthy-weight asthma, obesity alone and those that are healthy-weight without asthma. We identify that distinct CD4+T cell subtypes underlie CDC42 upregulation and Th1 polarization, but both are enriched for steroid resistance. NFκB was upregulated in additional clusters suggesting global changes in obese asthma CD4+T cells rather than any single cell subtype that may be targetable for therapeutics for obesity-related asthma. However, enrichment of distinct clusters of cells with specific genes that are known perpetrators of steroid resistance identify the potential to consider gene-targeted cellular therapies for obesity-related asthma.

While we report for the first time on distinct CD4+T cell subtypes associated with CDC42 upregulation, Th1 polarization, and steroid resistance in pediatric obese asthma, our study has limitations. There was lack of complete correlation between cell clusters and cell subtypes defined by Azimuth classification, both of which are identified by computational analysis of gene expression rather than protein expression. While this is in keeping with the pleiotropic nature of CD4+T cells, it limits our ability to use these observations to conduct targeted downstream functional analysis on a specific CD4+T cell. Given the cross-sectional nature of our investigation, we are unable to define which cell type initiated the observed cellular patterns. Few genes correlated with FEV_1_/FVC and only one correlated with ERV in obese asthma, an observation that differs from our prior findings.^13^ These divergent observations are likely influenced by differences in our cohorts, where the current one included children ages 7-18 years as compared to the prior one where the age range was 7-11 years, as well as the fact that the majority of the recruitment of the current cohort occurred during the pandemic when asthma was better controlled among children and adolescents. We therefore speculate that better disease control decreased the likelihood of large proportions of circulating cells to be in an activated state.^54^ This speculation is supported by the fact that clusters with CDC42 upregulation comprised of naïve and central memory cells that are quiescent as compared effector memory cells, but are poised to activate Th cells, an observation that also highlights the chronicity of the disease. Lastly, our cohort is comprised solely of underserved children. Although their ancestry and exposure to social determinants of health may underlie some of our observations that may limit the generalizability of our findings, investigations like ours on obese asthma in these populations are needed to identify the distinct cellular footprints unique to these populations that may explain their higher disease burden.

In conclusion, there is substantial evidence to support that obesity-related asthma has a non-atopic phenotype. Contrary to our hypotheses that a single cell cluster or a few clusters underlie the non-atopic phenotype, we found that the majority of CD4+T cells are qualitatively and quantitatively programmed differently in obese asthma as compared to healthy-weight asthma, obese alone and healthy-weight without asthma but have distinct cell subtypes enriched for CDC42 upregulation and Th1 polarization, that are both associated with steroid resistance. Conserved genes in these cell subtypes correlate with the obese asthma phenotype. These observations suggest that targeted identification of CD4+T cell by its gene profile is needed to further define the underlying pathways to target for therapeutics for obese asthma.

## Methods

### Study population

The study cohort comprised of 135 African American and Hispanic children, ages 7-18 years, including 54 children with obesity-related asthma, 27 with healthy-weight-asthma, 27 with obesity- alone, and 27 with healthy-weight without asthma (healthy-weight controls), that were recruited between June 2019 and June 2023. Obesity was defined as body mass index (BMI) >95^th^ percentile for age and sex.^55^ Asthma was defined based on documentation of physician diagnosis, reversible airflow obstruction, and ongoing asthma therapy in the electronic medical records.^56^ Children underwent anthropometric measurements, pulmonary function testing, and a fasting blood draw at the study visit.^12^ The study was approved by the Institutional Review Board at Children’s National Hospital and Albert Einstein College of Medicine, the two sites of study recruitment.

### CD4+ cell isolation

Peripheral blood mononuclear cells (PBMCs) separated from blood by Ficoll Hypaque method underwent CD4+T cell separation using magnetic bead-based negative isolation (StemCell Technologies, Tukwila, WA) to prevent *ex-vivo* stimulation.^12^

### Single-cell RNA Seq analysis

Six biological replicates of CD4+T cells from each study group underwent single-cell RNA-seq (sc-RNA-seq) on the 10X Genomics platform,^57, 58^ using the CITE-Seq approach that allowed cell surface labeling with antibody derived tags (ADTs), including CD4, CD25, CD127, CXCR3, CCR6, CCR10 and CCR4 (Biolegend, San Diego, US). Using Hash Tag Oligonucleotides (HTOs), 4 samples (one from each group) were multiplexed on one lane and sequenced on Novaseq 6000 (Novogene Corporation, Sacramento, CA).^59^

The sequencing results were run through the pipeline offered as part of *CellRanger* v7.1.0 (10X Genomics) for quality control (QC), barcode counting, and ADT quantification. *Seurat* R package v4.0 was used to demultiplex and assign samples to study groups, to classify cells into clusters, visualize using the UMAP feature, and calculate cluster-specific cell proportions and percent ADT expression.^60^ One sample in the healthy-weight control group failed QC so was excluded from the analysis. Cluster-specific conserved genes, comprised of genes enriched in a cluster in all samples compared to other clusters, were identified using *FindConservedMarkers* function within the *FindAllMarkers* function. Azimuth was applied for unbiased CD4+T cell subtype classification of clusters **[Table S1]**.^25^ To classify clusters into known Th cell subsets,^61^ we quantified cluster-specific gene expression of transcription factors (TFs), including *TBX21,* which encodes for the T-bet TF for Th1 cells, *GATA3,* which encodes for the GATA-3 TF for Th2 cells, *RORA* and *RORC,* which encode for the RORα and RORγt TFs for Th17 cells, and *FOXP3,* which encodes for the FoxP3 TF for T regulatory cells.^26^ Using canonical correlation analysis, we identified clusters in which *CDC42* was differentially expressed in obese asthma as compared to the three other groups and compared additional genes in the CDC42 pathway in these clusters.^12^ We also compared between-group expression of 50 genes curated from the literature for their association with Th1 responses, steroid resistance, and NLRP3 inflammasome **[Table S2]**. ^18, 20, 21, 22, 23, 24^ Conserved genes in clusters enriched for *CDC42* and Th1 responses were analyzed on NetworkAnalyst.^62^

### Bulk RNA-seq

Five hundred nanogram of RNA from CD4+T cells underwent bulk RNA-seq using KAPA stranded mRNA-Seq kit (KAPA Biosystems, Wilmington, MA). Transcripts were aligned to hg38 assembly using *STAR aligner* (v2.4.2a)^63^, annotated using GenCode (v19), and analyzed on DESeq2. Normalized gene counts were analyzed using linear regression modeling to elucidate between-group differences, adjusting for covariates associated with gene expression variance.^12^

### Immunoblot analysis

To verify differential expression of *CDC42* and elucidate its downstream signaling pathways, protein expression was quantified in 5 biological replicates of CD4+T cells from each study group that were distinct from the 6 replicates that underwent sc-RNA-Seq analysis. CD4+T Cells were stimulated with Dynabeads Human CD3/CD28 T-Activator (Thermo Fisher Scientific, MA) for 30 minutes and 2 hours. Four microgram protein per sample was resolved on a 4-15% gel, was electro-blotted to a membrane, and was probed first with CDC42, phospho- and total p70- Ribosomal protein S6 kinase (p70S6K1), phospho-p38 mitogen-activated protein kinase (p38MAPK), and α-tubulin (loading control) antibodies. The blot was then stripped and probed with phospho-NF-κB and total p38MAPK antibodies, and then stripped again and probed a third time with total NF-κB antibody. Phospho-p70S6K1, total NF-κB and α-tubulin antibodies were generated in mouse and others were generated in rabbit; the secondary anti-mouse antibody fluoresced at 680 nm (green) and anti-rabbit antibody fluoresced at 780 nm (red). Antibodies were purchased from Abcam, Cambridge, UK, or Cell Signaling, MA. Bands were quantified with LI-COR Odyssey System and images analyzed using ImageJ software.^64^ Analysis of variance on GraphPad Prism v.10 was used to quantify between-group differences in protein levels normalized to α-tubulin.

### Statistical Analysis

Analyses of sc-RNA-seq, bulk RNA-seq, and immunoblotting are described above. To elucidate clinical relevance of genes identified in sc-RNA-seq analysis, we used normalized gene counts from bulk-RNA-seq data and applied Pearson correlation tests to identify genes associated with FEV_1_/FVC ratio and percent predicted ERV, given the relevance of these two pulmonary function indices in obesity-related asthma.^11, 53^ We applied repeated permutation analysis (100 times) to identify random association of genes with FEV_1_/FVC ratio and percent predicted ERV; the proportion of genes associated with FEV_1_/FVC ratio or percent predicted ERV higher than random association was defined as measure of enrichment of association of gene expression with pulmonary function.

### Data and code sharing statement

Sequencing data will be made available in GEO NCBI database. R code used for data analysis will be shared upon request.

## Acknowledgements

We would like to acknowledge the children and their caregivers who participated for their contributions to this study.

## Author contributions

DT analyzed the transcriptomic data. YBW, SD, LC, and KC contributed to study participant recruitment and processing of blood samples. CY and AR conducted wet bench experiments. JMG and DR conceived the study and DR supervised the execution of the project. DR drafted the manuscript. DT, YWB, SD, AR, LC, KC, CY, and JMG reviewed, edited, and approved the manuscript. The authors have declared no conflict of interest.

